# Targeting von Willebrand factor selectively under inflammatory conditions

**DOI:** 10.1101/2025.08.13.670163

**Authors:** Gianluca Interlandi, Victoria S. Carter, Yi Wang, Alexander St. John, Jennie Le, Junmei Chen, José A. López, Xiaoyun Fu

## Abstract

It is challenging to develop anti-thrombotic therapeutics to treat or prevent pathological thrombus formation without increasing the risk of bleeding. Currently available anti-coagulant drugs render blood thinner irrespective of whether an inflammatory pro-thrombotic condition exists or whether clotting needs to occur because of traumatic vessel rupture. It is desirable to develop a drug that is active only under the oxidizing conditions present during inflammation. The blood protein von Willebrand factor (VWF) plays a key role in initiating blood clotting and it has been the target of anti-thrombotic therapies. It has been shown that oxidizing agents released during inflammation activate VWF by converting methionine residues within its domains to methionine sulfoxide. A previous study by us developed a method to computationally screen for drugs that inhibit VWF more strongly in the presence of oxidizing conditions. The computations suggested in particular a drug, lumacaftor, that could inhibit VWF function selectively in the presence of oxidized methionine residues. Here, we developed an enzyme-linked immunosorbent assay to test the effect of drugs on the ability of VWF to bind to the platelet surface receptor glycoprotein Ib*α* comparing oxidizing and non-oxidizing conditions. The results indicate that lumacaftor may indeed have the desired properties of inhibiting VWF selectively when oxidized. Because of its simplicity, the assay provides a high-throughput method to efficiently screen multiple drugs against VWF in both its oxidized and unoxidized state.

## Introduction

Currently available anti-thrombotic therapeutic drugs, often referred to as “blood thinners”, generally reduce the ability of blood to clot irrespective of whether clotting is due to pro-thrombotic conditions leading to pathological thrombus formation or to the physiological response to vessel rupture. In fact, commonly available anti-thrombotic drugs such as warfarin,^1^ caplacizumab,^2,3^ which targets VWF directly and is used to treat thrombotic thrombocytopenia purpura, or apixaban^4^ (sold as Eliquis), an inhibitor of factor Xa, all carry the risk of bleeding as side effects. Hence the question: Is it possible to design a therapeutic that blocks the formation of a pathological thrombus without affecting the beneficial haemostatic response of blood to clot in case of injury? Such a drug would need to distinguish between an inflammatory pro-thrombotic environment and a situation where clotting is activated to stop blood loss.

Inflammation has been linked to to an increased risk of thrombosis.^5^ During the inflammatory response, hydrogen peroxide is released, which is converted to hypochlorous acid (HOCl) through the action of myeloperoxidase. The oxidizing agent HOCl causes the conversion of methionine residues to methionine sufoxide modifying the structure and function of several blood proteins inducing a pro-thrombotic state.^6^ An example of a blood protein whose pro-thrombotic function has been shown to be increased following oxidation of its methionine residues is von Willebrand factor (VWF).^7^ Multiple studies have investigated VWF as a possible target of anti-thrombotic therapies^8–11^ since VWF is thought to have a central role in pathological thrombus formation.^12–14^ Hence, studying how VWF is activated under various conditions in particular in the presence of oxidants may provide a pathway how to design an inhibitor that targets VWF selectively under a pro-thrombotic inflammatory environment.

The protein VWF tethers blood platelets to the site of vascular injury in the early stages of haemostasis. It can be described as a relatively long multimeric chain where monomers are linked to each other through disulfide bonds at the N- and C-terminii^15^ (Figure 1). Each monomer consists of a number of domains that are covalently linked to each other^15^ (Figure 1). The A1 domain is responsible for mediating the platelet-binding function of VWF by binding to glycoprotein Ib*α* (GpIb*α*) located on the surface of blood platelets. It has been shown that the function of the A1 domain can be inhibited by its neighboring domains, specifically, the D’D3 domains located N-terminally^16,17^ and the A2 and A3 domains located C-terminally^18^ (Figure 1). Normally, tensile force generated by shear stress present in flowing blood separates neighboring domains from A1, which becomes exposed and hence is able to bind GpIb*α*.^19^ Recently, oxidizing conditions have been shown to increase the platelet-binding function of VWF and this has been linked to the oxidation of methionine residues in the A1, A2 and A3 domains.^7^ Furthermore, molecular dynamics (MD) studies have provided evidence that oxidation of methionine residues destabilizes the fold of the A2 domain^20^ and disrupts the interface between the A1 and A2 domains.^21^ The separation of the A1 and A2 domains from each other due to methionine oxidation is consistent with experimental evidence that oxidation increases the GpIb*α*-binding function of a A1A2A3 domains construct but not of isolated A1.^21^ Hence, studying how methionine oxidation alters the interfaces between A1 and the neighboring domains, which normally inhibit its function, can lead to the discovery of therapeutics that block VWF function only under inflammation-induced oxidizing conditions while preserving the haemostatic function of VWF.

**Figure 1:**
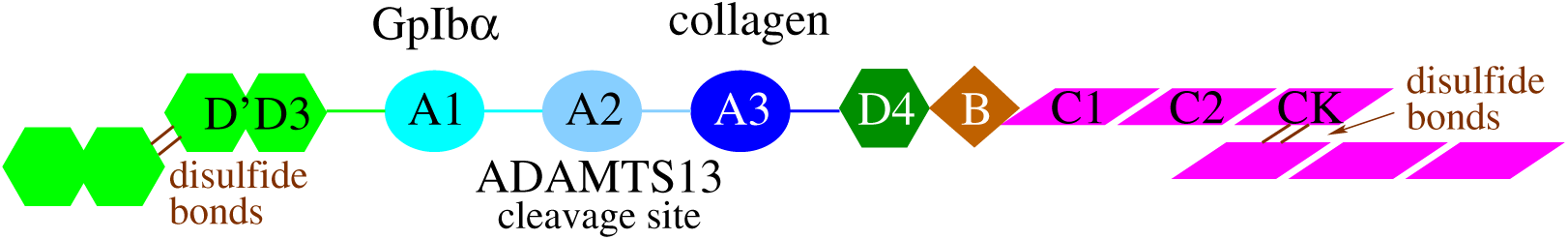
Overview of VWF structure. Schematic representation of a VWF monomer. Indicated are the substrates of the A1 and A3 domain, and the fact that the A2 domain contains the cleavage site for ADAMTS13 (Figure from reference^21^).

To achieve this, it is necessary to develop tools to efficiently screen for drugs that compensate the activating effect of oxidation on VWF while leaving the platelet-binding function of unoxidized VWF unaltered. This would be useful for example to search for drugs that are already available or even already approved by the Food and Drug Administration (FDA) and could be repurposed as anti-thrombotic therapeutics if they have the desired effect on VWF. A previous study by us used a combination of molecular docking, molecular dynamics (MD) simulations and free energy perturbation (FEP) calculations to search for drugs in the ZINC15 database^22^ that are more likely to bind at the A1-A2 interface in the presence of oxidized methionine residues.^23^ Such a drug would act as a “glue” between the two domains hence restoring the inhibitory function that A2 normally has on the A1 domain. The search was restricted to drugs that are already FDA approved and easily available. The study suggested two drugs, budesonide and lumacaftor, that may target VWF under oxidizing conditions.^23^ In particular, lumacaftor was found *in silico* to be more likely to bind at the A1-A2 interface under oxidizing than under normal conditions.^23^ Hence, lumacaftor may be a candidate for a drug that prevents thrombosis while maintaining haemostasis. Available compound databases can contain millions of entries and screening them *in vitro* may not be feasible. The computational method described in^23^ can be applied to narrow down the number of molecules to be tested *in vitro* from millions to possibly thousands. However, a high-throughput experimental method is still necessary to efficiently test such a large number of molecules in a cost-effective manner.

Here, we present a method based on an enzyme-linked immunosorbent assay (ELISA) to measure the binding activity of VWF to GpIb*α* comparing oxidized and unoxidized VWF in the absence and the presence of lumacaftor or budesonide in order to test the previously obtained computational predictions. Furthermore, mass spectrometry was used to ascertain whether the increased VWF activity with oxidation was correlated to the rate of oxidation of methionine residues. The advantage of ELISA is that it allows comparing multiple conditions at the same time as each plate used here contains 96 wells, and more conditions can be compared with plates containing an even larger number of wells. Hence, the method presented here can serve as the basis for a high-throughput screening assay. The search can then be expanded to a larger number of drugs, which can be tested through the combination of computational predictions described earlier^23^ and the experimental assay developed here.

## Materials and Methods

### Reagents for ELISA

Purified full-length human von Willebrand factor was purchased from Prolytix (HCVWF-0190). Recombinant human GpIb*α* containing a histidine tag at the C-terminus was purchased from bio-techne (4067-GP). Generally, stock solutions were prepared using 0.2% BSA-PBS as the buffer. The ELISA was performed using Immulon 4 HBX 96-well plates.

### ELISA experiments

Wells of a 96-well plate were coated each with 100 *µ*l of a solution containing 1 µg of VWF. Subsequently, the plate was incubated at 37^◦^ C for 90 minutes, after which the coated wells were washed three times with PBS as after each incubation step. Assuming a molecular weight of a VWF monomer to be approximately 260 kDa, the VWF solution used here corresponds to a concentration of about 38.5 nM of VWF monomers. It is important to note that VWF is multimeric, but each monomer contains one A1 domain, i.e., one binding site for GpIb*α*. The protein VWF was either left unoxidized or was oxidized with HOCl. Oxidation of VWF was either performed in-vial before adding the solution to the wells, or in-plate after VWF had been adsorbed on the plate. To perform in-vial oxidation, HOCl was added to the VWF solution to achieve the final concentration indicated in the experiment and the vial was rotated at 37^◦^ C for 60 minutes, followed by quenching with excess free methionine rotating at 37^◦^ C for 15 minutes. In-plate oxidation was performed by adding 100 *µ*l of HOCl at the concentration indicated in the experiment and incubating for 60 minutes at 37^◦^ C. This was followed by quenching with excess free methionine while incubating for 15 minutes. To block the wells, 1% BSA-PBS was added and the plate was incubated overnight at 4 ^◦^ C.

To test the effect of drug molecules, a 100 *µ*l solution containing a drug at the indicated concentration was added to specific wells and incubated at 37^◦^ C for 60 minutes. Finally, recombinant GpIb*α* was added to all wells at a concentration of either 1-fold, two-fold or five-fold the concentration of the VWF solution used in the first incubation step (38.5 nM). The different concentrations of GpIb*α* are labeled in Section 3 as “1x GpIb*α*”, “2x GpIb*α*” and “5x GpIb*α*”, respectively. Then, a horseradish-conjugated anti-histidine antibody (BioLegend Clone J099B12) was added to the wells and incubated at 37^◦^ C for 60 minutes. Subsequently, TMB was added, the reaction was stopped with a stop reagent (Sigma-Aldrich S5814) and the plate was read at 450 nm.

### Nano flow liquid chromatography tandem mass spectrometry

Oxidized and control unoxidized human VWF samples were reduced with dithiothreitol (Bio-RAD), alkylated with iodoacetamide (Bio-RAD), and digested with Trypsin/LysC (Promega). The resulting tryptic peptides were desalted using a C18 cartridge (3M Science) and analyzed by nano flow liquid chromatography tandem mass spectrometry (nanoLC-MS/MS) using a Thermo Scientific Orbitrap Fusion^TM^ Lumos^TM^ Tribrid mass spectrometer coupled with a Waters nanoACQUITY Ultra Performance LC system. Peptides were separated at a flow rate of 300 nL/min on an ACQUITY UPLC M-Class HSS T3 Column (100 x 0.075 mm, 1.8 *µ*m, Waters), using solvent A (0.1% formic acid in water) and solvent B (0.1% formic acid in acetonitrile). Peptides were eluted with a linear gradient: 5%-25% solvent B over 38 min, 25%-90% solvent B over 5 min, and 90%-95% solvent B over 15 min. Data acquisition was performed in positive ion mode with a Parallel Reaction Monitoring (PRM) method targeting predefined peptides. Peptide peak areas were quantified using Thermo Scientific^TM^ Xcalibur^TM^ software (v2.2). The percent oxidation of individual methionine residues was calculated by dividing the peak area of the methionine-containing oxidized peptide by the sum of the peak areas of both oxidized and unoxidized peptides.

### Obtaining platelet-poor plasma from trauma patients

This collection protocol was approved by the University of Washington Human Subjects Division Institutional Review Board (protocol number STUDY00008295).

Blood samples were collected from severely injured patients soon after injury. Adult trauma patients at least 18 years old were included if they met institutional criteria for full trauma team activation at Harborview Medical Center, Seattle, Washington, USA. These criteria use mechanism of injury and early physical exam findings to identify patients at high risk for severe injury. Patients were excluded if they arrived at the Emergency Department more than 3 hours after injury or if they received more than 2 liter of crystalloid fluid, 1 unit of whole blood, or 2 units of component blood products of any kind prior to blood sampling. Patients were also excluded if they suffered burns of more than 5% total body surface area, received tranexamic acid (TXA) prior to initial blood draw, were known to have congenital coagulopathy or take anticoagulants or antiplatelet medication, or were pregnant, prisoners, or transferred from another facility.

Due to minimal risk to the patient, the requirement for consent was waived. However, patients were allowed to opt out of the study at any time during enrollment or afterward. After collection of the samples into citrated vacutainers, the blood was centrifuged to create platelet-poor plasma (1,200 g for 15 min), and the plasma was aliquoted and frozen. Plasma samples were collected from a total of 29 patients, and VWF antigen levels were determined for each patient. For this study, we used the plasma sample with the highest concentration of VWF, which was 22.05 *µ*g/ml.

### VWF antigen level

VWF antigen levels were quantified using ELISA. A 96-well plate was coated with an anti-human VWF antibody (DAKO #A0082). The surface was blocked with 5% BSA in PBS for 1 hour at room temperature. Patient plasma was added to the wells, incubated for 1 hour and then washed. A second, detecting antibody, anti-human VWF conjugated with horseradish peroxidase (DAKO #P0226), was added and the plate was incubated for 1 hour. After washing, TMB was added and the plate was read in a plate reader. The concentration of VWF was calculated by using a regression line equation obtained with reference plasma.

### Ristocetin induced platelet agglutination

Ristocetin induced platelet agglutination (RIPA)^24^ was performed using the AggRAM Analyzer (Helena Laboratories). Standardized human platelets (Bio/Data Corporation) were mixed with ristocetin with a final concentration (after adding all reagents) of 0.55 mg/ml. Platelet-poor plasma either derived from a trauma patient or normal pooled plasma (Precision Biologics) was then added to the platelet/ristocetin mixture and the platelet agglutination reaction was monitored for five minutes. Both, plasma from the trauma patient and normal pooled plasma, were either pretreated with lumacaftor (final concentrations of either 10 or 100 *µ*M) or left unaltered for comparison. All plasma samples, whether pretreated with lumacaftor or left unaltered, were incubated at room temperature for one hour while being rotated at 24 rpm in a Mini LabRoller™(Labnet).

## Results

### Activation of VWF correlates with methionine oxidation

In order to design drugs that target VWF under oxidizing conditions, it is necessary to evaluate whether the increased activity of VWF in the presence of HOCl is linked to the increased rate of methionine oxidation. Establishing such a link makes it possible to use structure-based drug design to narrow down from a large database a set of molecules to be tested *in vitro* as described in a previous publication.^23^ Increased platelet-binding activity of VWF due to oxidation through HOCl has been previously reported using a ristocetin-induced platelet agglutination (RIPA) assay.^7^ However, ELISA is a much more convenient assay to test the activity of VWF under various conditions because it allows measuring multiple combinations of oxidant and drug concentrations in one single plate. Hence, we tested whether 1) the oxidation-induced activation of VWF observed in RIPA^7^ can be recapitulated in a ELISA, and 2) the increased activation is related to the oxidation rate of methionine residues.

In order to facilitate nanoLC-MS/MS analysis of the methionine oxidation rate in VWF, oxidation was performed in-vial (See Section 2). We observed that increasing concentrations of HOCl increased the activity of VWF in binding GpIb*α* (Figure 2A). Furthermore, increasing concentrations of HOCl also caused increasing rates of oxidation of specific methionines for which it was possible to determine the oxidation state in our nanoLC-MS/MS assay (Figure 2B). Hence, increasing concentrations of HOCl cause both, an increase in binding activity and a higher rate of methionine oxidation.

**Figure 2:**
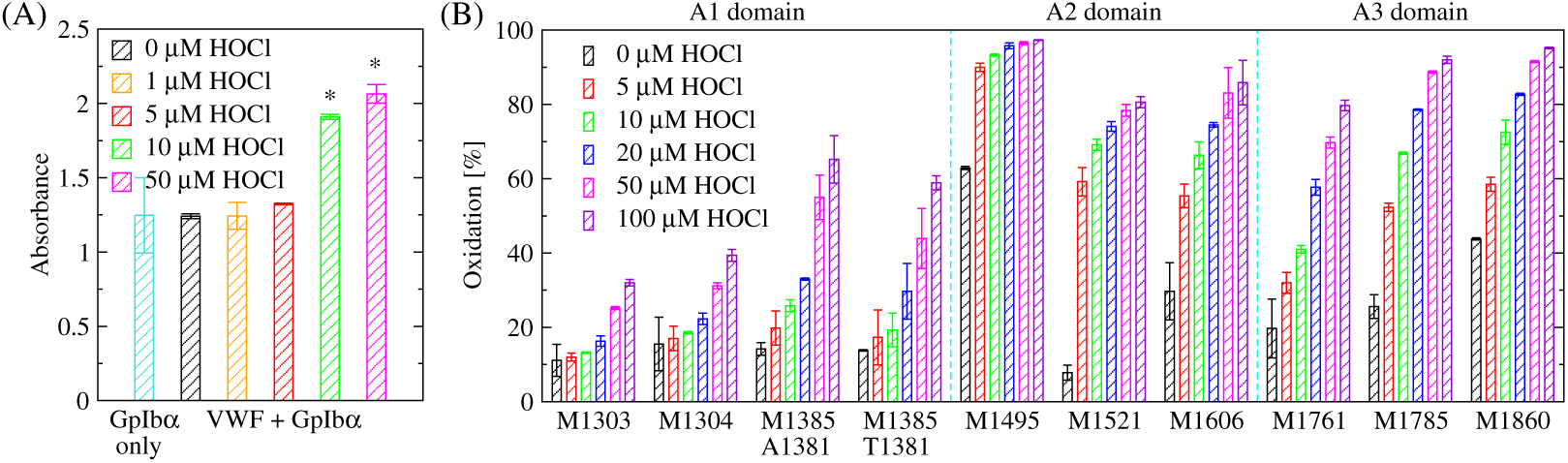
Titration with oxidation performed in-vial. (A) Increase in VWF binding activity with increasing concentration of HOCl in ELISA. The binding experiments were performed with a 5x GpIb*α* concentration (see Section 2). A “+” indicates a difference that is marginally statistically significant (p-value between 0.05 and 0.1) while a “*” denotes a statistically significant difference (p-value < 0.05) when comparing oxidized to non-oxidized (no ox) VWF. “GpIb*α* only” indicates that no VWF was used. (B) nanoLC-MS/MS of methionine residues as a function of HOCl concentration. The values for M1385 are reported for variants containing alanine or threonine at position 1381 since VWF found in humans consists of a population with both variants. Vertical dashed cyan lines separate residues located within A1, A2 and A3 domains.

In order to test whether oxidation increases the activity of VWF also when HOCl is added after VWF is adsorbed to the plate, we performed an ELISA where various concentrations of HOCl were added in-plate after incubating overnight with 1% BSA-PBS. We observed that VWF binding to GpIb*α* increased with increasing oxidant concentration where 50 *µ*M and 100 *µ*M HOCl produced results that were significantly different from the activity of non-oxidized VWF (Figure 3). Since we have observed more consistent outcomes with in-plate rather than in-vial oxidation, the following ELISA experiments were performed with in-plate oxidation.

**Figure 3:**
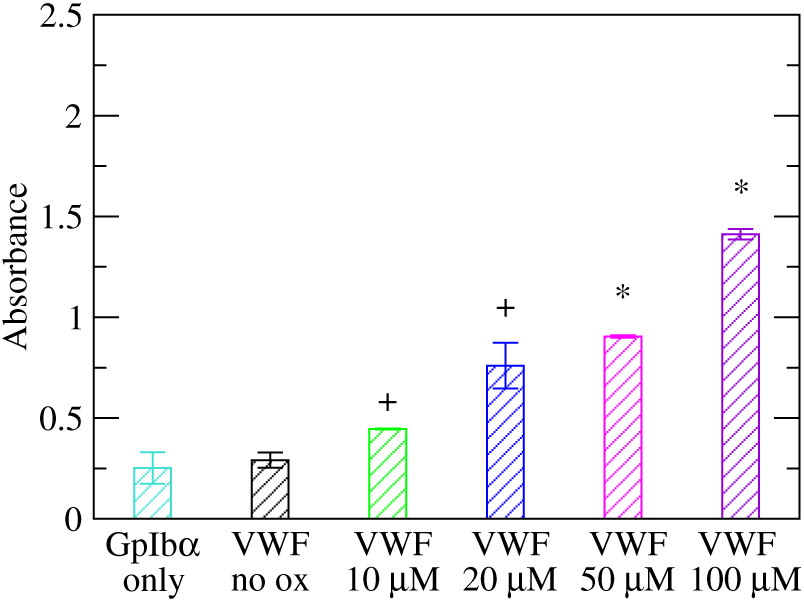
Titration with oxidation performed in-plate. Increase in VWF binding activity with increasing concentration of HOCl in ELISA. The binding experiments were performed with a 2x GpIb*α* concentration (see Section 2). A “+” indicates a difference that is marginally statistically significant (p-value between 0.05 and 0.1) while a “*” denotes a statistically significant difference (p-value < 0.05) when comparing oxidized to non-oxidized (no ox) VWF.

### ELISA tests of lumacaftor and budesonide

In a previous computational study, we identified two FDA approved drugs, lumacaftor and budesonide, that bind at the interface between the A1 and A2 domains of VWF in the presence of oxidized methionine residues^23^ suggesting that such drugs could restore the inhibitory function of the A2 domain on A1, the domain responsible for binding to the platelet surface receptor GpIb*α*. In particular, lumacaftor was found to bind at the A1-A2 domains interface significantly stronger under oxidizing than normal conditions^23^ suggesting that lumacaftor could be a candidate to inhibit VWF selectively under inflammatory conditions. These drugs were tested here through the ELISA described in Section 2.

Different concentrations of lumacaftor were added to the plate after it was either left unoxidized or oxidized with various amounts of HOCl. While oxidation increased the binding activity of VWF, lumacaftor decreased it significantly (Figure 4A). Oxidation was observed to also increase the background non-specific adhesion of GpIb*α* to the plate although with a significantly smaller effect than in the presence of adsorbed VWF. In fact, subtracting the background for each respective HOCl concentration still yielded statistically significant differences when comparing oxidized to unoxidized VWF and when comparing with and without the addition of lumacaftor to oxidized VWF (Figure 4B). Furthermore, even in the absence of oxidant, wells with VWF had a significantly larger amount of bound GpIb*α* than wells containing only blocking buffer (Figure 4A). This observation indicates that it is possible to measure the activity of unoxidized VWF with this assay, which allows testing the effect of a drug also on unoxidized VWF, which is normally less active. Notably, lumacaftor did not alter the function of unoxidized VWF (Figure 4). This is desirable for a drug meant to prevent thrombosis while maintaining haemostasis.

**Figure 4:**
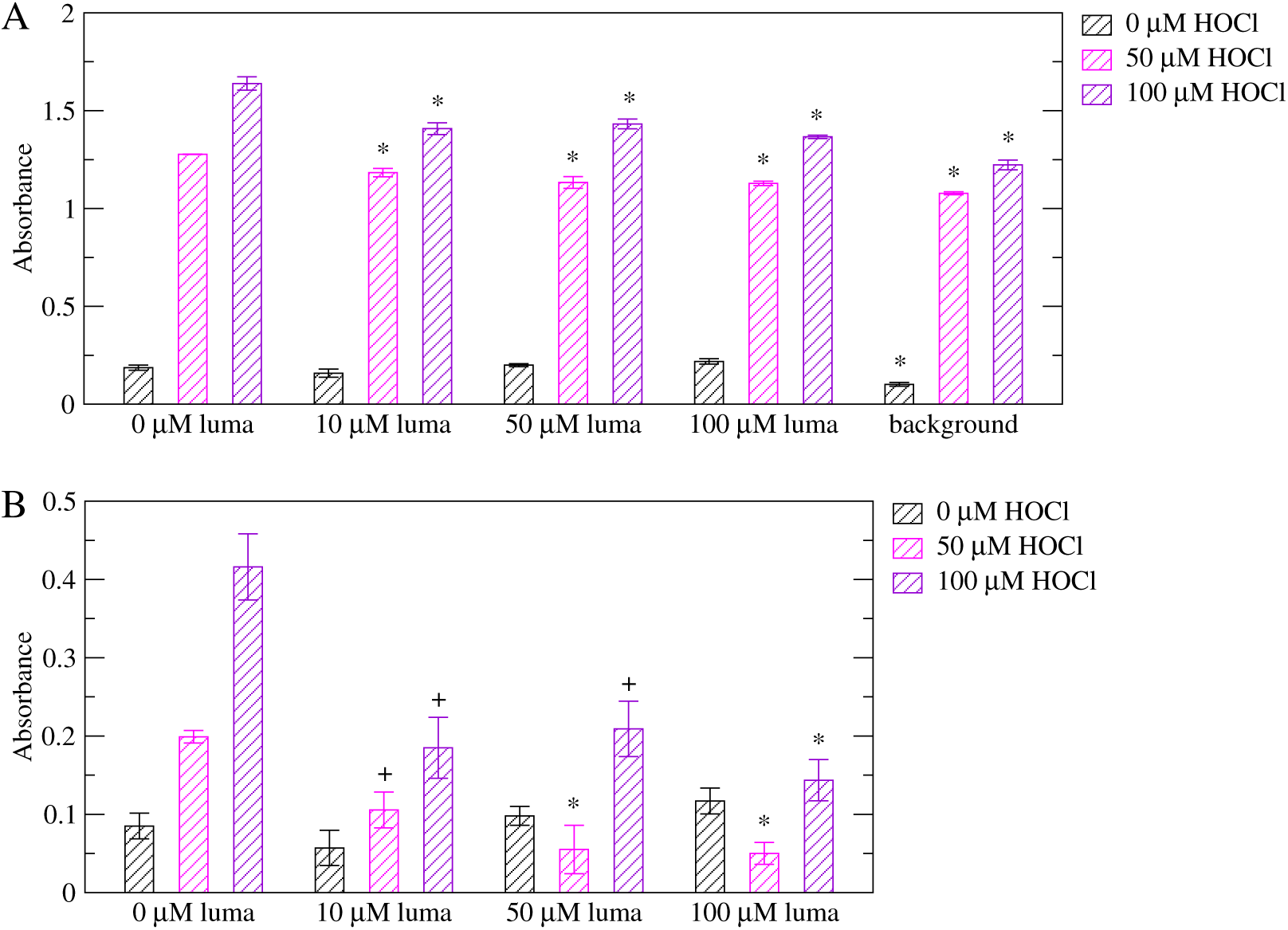
ELISA testing the effect of lumacaftor on VWF activity. (A) Comparison of the binding activity of non-oxidized (0 *µ*M HOCl) and oxidized (50, 100 *µ*M HOCl) VWF in the absence and presence of lumacaftor. Experiments were performed with a 2x GpIb*α* concentration (see Section 2). A “*” denotes a statistically significant difference (p-value < 0.05) when comparing the reported value to wells not treated with lumacaftor while a “+” denotes a marginally statistically significant difference (p-value between 0.05 and 0.1). “luma” stands for lumacaftor while “background” indicates wells treated only with blocking buffer but no VWF. Reported are averages over two measurements while error bars denote standard errors of the mean (SEM). (B) The same as above, but the background is subtracted for each respective concentration of HOCl. Since the reported values are calculated by subtracting two measured values, error bars are calculated using the error propagation formula: 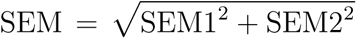 where SEM1 and SEM2 are the standard errors of the mean of the corresponding measurement with VWF and the background, respectively, as reported in panel (A).

The inhibitory function of lumacaftor on oxidized VWF was observed also when using a lower concentration of GpIb*α* (Figure 5), i.e., using 1x instead of 2x GpIb*α* (see section 2), in particular at 100 *µ*M of HOCl. Notably, using a lower concentration of GpIb*α* decreased the background noise under oxidizing conditions. In fact, with 2x GpIb*α*, the background signal was 84% ± 1% and 75% ± 2% of the signals from wells with VWF but no lumacaftor at 50 and 100 *µ*M HOCl, respectively. Using 1x GpIb*α*, the ratios of background to VWF decreased to 65% ± 7% and 59% ± 5% at 50 and 100 *µ*M HOCl, respectively. Hence, a lower concentration of GpIb*α* may improve the signal to background ratio.

**Figure 5:**
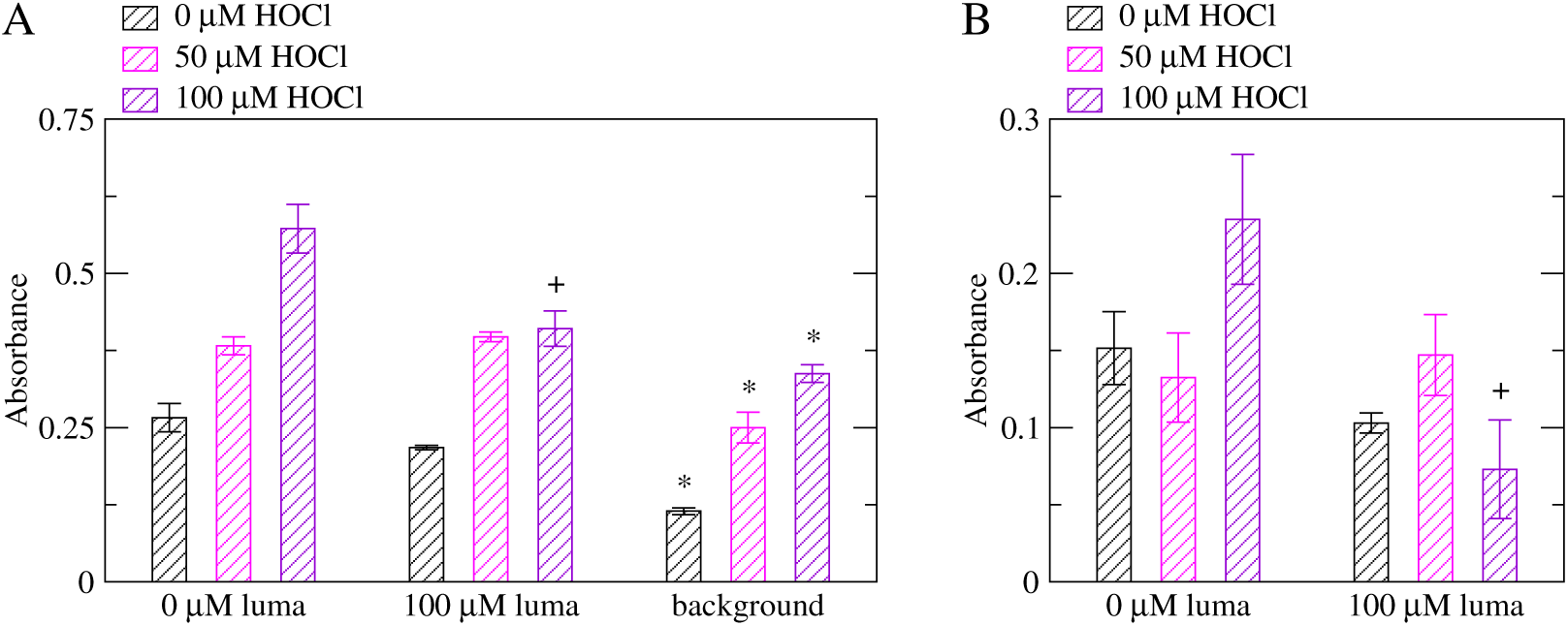
ELISA testing the effect of lumacaftor on VWF activity using a lower concentration of GpIb*α*. (A) Comparison of the binding activity of non-oxidized (0 *µ*M HOCl) and oxidized (50, 100 *µ*M HOCl) VWF in the absence and presence of lumacaftor. Experiments were performed with a 1x GpIb*α* concentration (see Section 2). A “*” denotes a statistically significant difference (p-value < 0.05) when comparing the reported value to wells not treated with lumacaftor while a “+” denotes a marginally statistically significant difference (p-value between 0.05 and 0.1). “luma” stands for lumacaftor while “background” indicates wells treated only with blocking buffer but no VWF. Reported are averages over two measurements while error bars denote standard errors of the mean (SEM). (B) The same as above, but the background is subtracted for each respective concentration of HOCl. Since the reported values are calculated by subtracting two measured values, error bars are calculated using the error propagation formula: 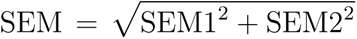 where SEM1 and SEM2 are the standard errors of the mean of the corresponding measurement with VWF and the background, respectively, as reported in panel (A).

According to our *in silico* study comparing the effects of various drugs on VWF, budesonide is probably able to bind to the A1-A2 domains interface but it may not be sensitive to the oxidation state of methionine residues.^23^ Hence, we tested budesonide with the ELISA assay developed here. While budesonide slightly decreased the GpIb*α*-binding activity of VWF under oxidizing conditions (not statistically significant in Figure 6A,B and marginally statistically significant in Figure 6C,D), the activity of unoxidized VWF was observed to be increased by the addition of budesonide (Figure 6). The discrepancy between the simulation predictions and the observations made with ELISA may be due to a different binding mode of budesonide to VWF that was not observed in the *in silico* study including that there could be a different way of modulating VWF actibity other than binding at the A1-A2 interface. Nonetheless, an increased binding activity under non-oxidizing conditions is not desirable for an anti-thrombotic drug.

**Figure 6:**
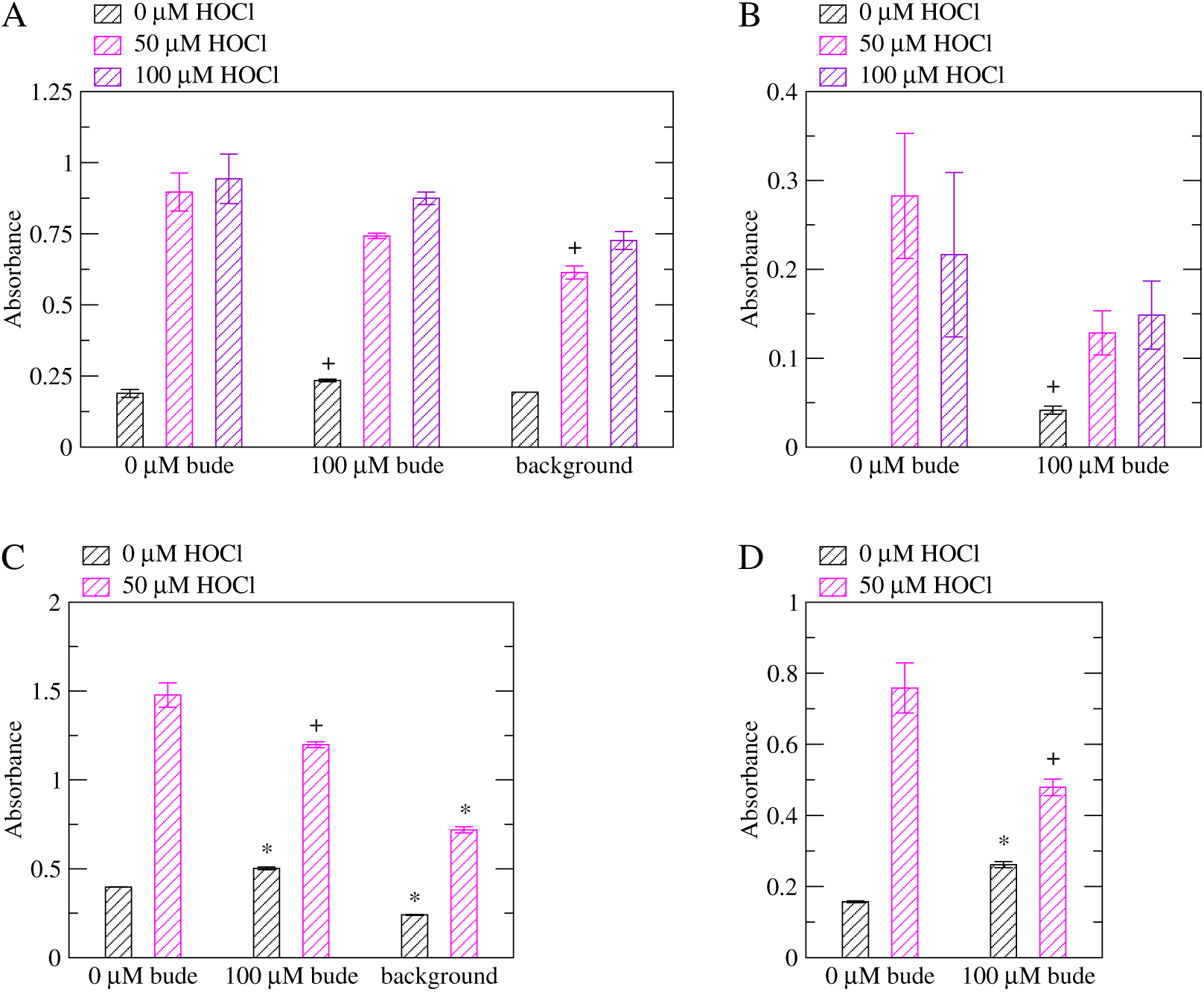
ELISA testing the effect of budesonide on VWF activity. (A) Comparison of the binding activity of non-oxidized (0 *µ*M HOCl) and oxidized (50, 100 *µ*M HOCl) VWF in the absence and presence of budesonide. Experiments were performed with a 1x GpIb*α* concentration (see Section 2). A “*” denotes a statistically significant difference (p-value < 0.05) when comparing the reported value to wells not treated with budesonide while a “+” denotes a marginally statistically significant difference (p-value between 0.05 and 0.1). “bude” stands for budesonide while “background” indicates wells treated only with blocking buffer but no VWF. Reported are averages over two measurements while error bars denote standard errors of the mean (SEM). (B) The same as above, but the background is subtracted for each respective concentration of HOCl. Since the reported values are calculated by subtracting two measured values, error bars are calculated using the error propagation formula: 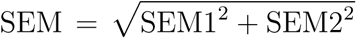 where SEM1 and SEM2 are the standard errors of the mean of the corresponding measurement with VWF and the background, respectively, as reported in panel (A). (C)-(D) Replicas of (A)-(B) but only with 0 and 50 *µ*M HOCl.

### RIPA assay with patient plasma testing lumacaftor

A blood sample was obtained from a trauma patient and platelets were removed to obtain platelet-poor plasma. The concentration of VWF was determined to be 22.05 *µ*g/ml through an antigen assay (see Materials and Methods). Such concentration is considered to be particularly elevated as the average concentration of VWF in the general population is reported to be 10 *µ*g/ml.^25^ As a comparison, the VWF concentration measured for the normal pooled plasma used as reference here was also 10 *µ*g/ml. We performed a RIPA assay^24^ (see Materials and Methods) to compare the VWF-induced platelet agglutination properties of patient plasma to normal pooled plasma in the absence and the presence of lumacaftor. We used two concentrations of lumacaftor, 10 *µ*M and 100 *µ*M. Platelet agglutination was monitored during five minutes after mixing standardized platelets with plasma samples in the presence of 0.55 mg/ml ristocetin (Figure 7A,C). We observed two interesting results. One is that platelet agglutination in the patient plasma was up to twice as strong as in the normal pooled plasma (Figure 7D).

**Figure 7:**
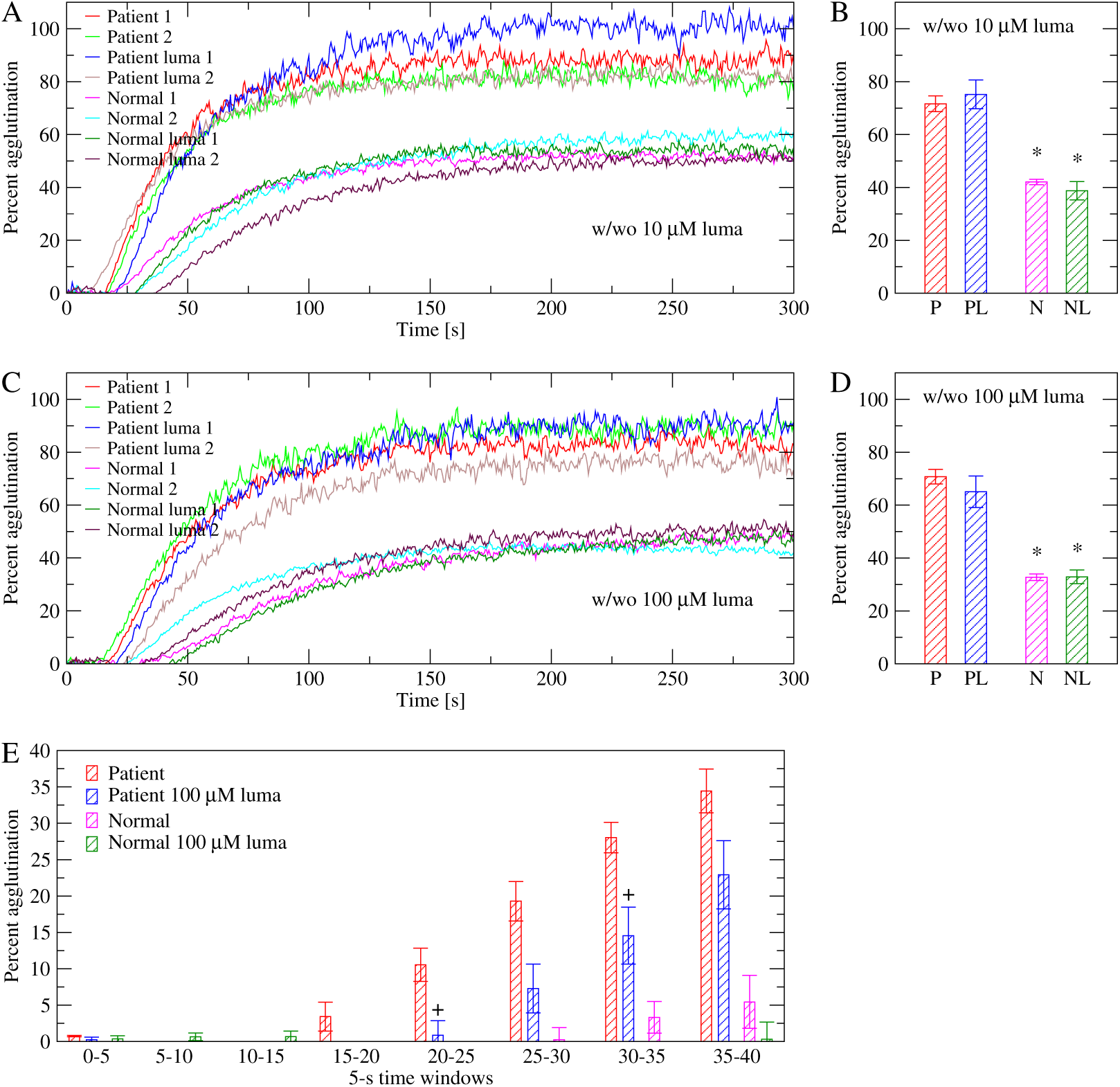
RIPA comparing patient to normal pooled plasma with and without lumacaftor. (A) Time series of platelet agglutination in the presence of 0.55 mg/ml ristocetin using plasma from a trauma patient and normal pooled plasma in the absence and presence of 10 *µ*M lumacaftor (abbreviated as “luma” in the figure). (B) Averages over the entire 300 seconds of monitoring for each condition measured in panel A. Error bars indicate standard errors of the mean (N=2). The asterisks indicate that the difference between patient and normal plasma (with or without lumacaftor) is statistically significant (p-value < 0.05 from a two-tailed Student’s t-test). (C) and (D) Same as in panels A and B, respectively, except that the concentration of lumacaftor used was 100 *µ*M. (E) Averages over 5-second windows for the first 40 seconds for the conditions measured in panel C. Error bars indicate standard errors of the mean (N=2). The “+” symbols indicate that the difference between patient plasma treated and untreated with lumacaftor is marginally statistically significant for that particular time window (p-value between 0.05 and 0.1 from a two-tailed Student’s t-test).

The other observation concerns the effect of lumacaftor on both patient and normal pooled plasma. Using 10 *µ*M of lumacaftor, there was no statistically significant difference between plasma pretreated with the drug and plasma left unaltered (Figure 7B). When using 100 *µ*M of lumacaftor, there was on average (calculated over the entire course of five minutes) a decrease in agglutination, but it was not statistically significant with either patient of normal pooled plasma (p-value larger than 0.1, Figure 7D). However, when averages were calculated over 5-second windows, we observed that the difference between treatment and no treatment was marginally statistically significant (p-value between 0.05 and 0.1) at two time windows during the first 35 seconds of the agglutination process when lumacaftor was added to the patient sample (Figure 7E). No difference was observed when lumacaftor was added to normal pooled plasma (Figure 7E).

## Discussion

It is desirable to discover a therapeutic that prevents pathological thrombus formation while allowing the haemostatic response in case of acute injury. Due to its central role in initiating the blood clotting process, the protein VWF serves as a key target for anti-thrombotic interventions. Oxidizing agents released during a pro-thrombotic inflammatory state have been shown to increase the platelet-binding function of VWF and to convert methionine residues to methionine sulfoxide.^7^ Hence, the question arises whether it is possible to find a drug that is sensitive to the oxidation state of VWF, inhibiting it only under the conditions that cause a pro-thrombotic environment.

Previous studies by us have suggested that oxidation of methionine residues removes the inhibitory function that the A2 domain normally has on A1.^20,21^ In particular, methionine residues located at the A1-A2 domains interface are converted to methionine sulfoxide in the presence of oxidizing agents^7^ disrupting the interaction between A1 and A2 domains.^21^ Hence, it is plausible that designing a drug that binds to oxidized methionine residues at the A1-A2 interface may act as a glue between the two domains and restore the inhibitory function of the A2 domain on the A1 domain. A computational study by us investigated such a question using a combination of molecular docking, molecular dynamics simulations and free energy perturbation calculations.^23^ It identified two drugs, budesonide and lumacaftor, that bound to both A1 and A2 domains in the presence of oxidized methionine residues.^23^ In particular, lumacaftor was found to bind to the A1-A2 interface significantly stronger when M1495 in the A2 domain was in its oxidized state.^23^ In this study, an assay based on ELISA was developed to test the effect of drugs such as budesonide and lumacaftor on the function of non-oxidized and oxidized VWF. nanoLC-MS/MS analysis revealed that the increase in VWF activity due to the addition of HOCl in the ELISA correlated with an increase in the rate of methionine oxidation (Figure 2) indicating that such an assay can be used to test the activity of VWF under oxidizing conditions. Computational high-throughput screening can be helpful to pre-select possible candidates from a very large database of molecules that have a particular effect on a protein. However, an efficient *in vitro* assay is required to rapidly screen the computationally pre-selected drugs on the function of a protein under various conditions such as in this case non-oxidized and oxidized VWF. The ELISA-based method presented here allows simultaneously testing multiple drugs under various concentrations comparing non-oxidized and oxidized VWF. Hence, it can serve as a high-throughput *in vitro* screening assay for drugs. The method was employed to test the effect of lumacaftor and budesonide on VWF. Analysis of the results indicated that lumacaftor reduced the function of oxidized VWF while leaving VWF function unaltered under normal conditions (Figures 4-5). On the other hand, budesonide had a limited effect on oxidized VWF and a slightly activating effect when VWF was left unoxidized (Figure 6).

Three conclusions emerge from this study. The first conclusion is that it is possible to design an ELISA to test the binding activity of VWF to the platelet surface receptor GpIb*α*. Previous studies have used a dynamic flow assay to measure the activity of full-length VWF or recombinant VWF constructs,^21,26^ which more closely mimics the effects of tensile force due to flowing blood. However, as shown previously^7^ and here (Figures 2 and 3), oxidation is also capable of activating VWF so that its GpIb*α* binding properties can be measured in a static assay. Furthermore, the ELISA was also able to detect binding of GpIb*α* to VWF not treated with HOCl (Figures 4-5) making it possible to test whether a drug alters VWF function under both oxidizing and non-oxidizing conditions. Such an ELISA-based approach allows for a rapid screening of different drugs under different conditions because such a static assay is simpler, less time intensive and less costly to setup then a dynamic flow assay as used in previous studies.^21,26^

The second conclusion, which is derived from the first, is that VWF can be activated also through adsorption on a polysterene surface as observed in the ELISA assay (Figure 4) in the absence of any modulator such as tensile force, oxidation or the drug ristocetin, which is known to be a potent activator of VWF.^27^ In a previous study that investigated how different material surfaces affect the GpIb*α*-binding function of the VWF A1 domain, polysterene and tissue-culture polysterene were found to increase the activity of A1 compared to glass.^28^ However, there are no studies reported to date how different material surfaces affect the function of full-length VWF. In this study, we observe that polysterene can activate VWF. Hence, similar studies testing the effects of various surfaces may provide useful for the design of body implants that do not induce pathological blood clots.

The third conclusion is that lumacaftor may be able to reduce the binding activity of VWF when oxidized (Figures 4-5). Furthermore, the function of unoxidized VWF was unaltered by the addition of lumacaftor (Figures 4-5) highlighting that this drug selectively targets VWF under oxidizing conditions. Hence, lumacaftor can be used as a framework to design an efficient and potent inhibitor of VWF that works selectively in the presence of inflammation-induced oxidizing agents. On the other hand, although the addition of budesonide slightly decreased the activity of oxidized VWF, it caused a slight increase of activity for unoxidized VWF (Figure 6). Hence, budesonide may not be as suitable as lumacaftor as the basis for the design of anti-thrombotic drug.

In conclusion, we have developed an ELISA-based assay to rapidly screen for drugs that decrease the activity of oxidized VWF while leaving the function of non-oxidized VWF unaltered. We were also able to measure binding of GpIb*α* to VWF adsorbed on the surface of a polysterene 96-well plate. This is novel because normally a modulator such as ristocetin or shear stress has been used to detect binding of GpIb*α* or platelets to VWF. This study is also an example of how computational docking and MD simulations as performed in a previous study by us^23^ can be combined with an *in vitro* assay to efficiently screen a library of drug molecules. Furthermore, repurposing already FDA approved drugs can greatly simplify the process to find a better therapeutic for a particular use. In this case, the goal is to target VWF selectively under oxidizing conditions in order to treat or prevent thrombosis while allowing the natural haemostatic response.

## Acknowledgments

This research was financially supported by NIH grant R01HL153253 and a Phase 1 Technology Commercialization Award from the Washington Research Foundation. The trauma patient samples were collected under the support of NIH grant K08HL146840.

